# Early founder effects have determined paternal population structure in the Faroe Islands

**DOI:** 10.1101/2024.08.27.601563

**Authors:** Allison E. Mann, Eyðfinn Magnussen, Christopher R. Tillquist

**Author notes:** These authors contributed equally to the project (Allison E. Mann, Christopher R. Tillquist). Correspondence Christopher R. Tillquist.

## Abstract

The Faroe Islands are a small archipelago located in the North Atlantic likely colonized by a small group of founders sometime between 50 and 300 CE. Post colonization, the Faroese people have been largely isolated from admixture with mainland and other island populations in the region. As such, the initial founder effect and subsequent genetic drift are likely major contributors to the modern genetic diversity found among the Faroese. In this study, we assess the utility of Y-chromosomal microsatellites to detect founder effect in the Faroe Islands through the construction of haplotype networks and a novel empirical method, mutational distance from modal haplotype histograms (MDM), for the visualization and evaluation of population bottlenecks. We compared samples from the Faroe Islands and Iceland to possible regional source populations and documented a loss of diversity associated with founder events. Additionally, within-haplogroup diversity statistics reveals lower haplotype diversity and richness within both the Faroe Islands and Iceland, consistent with a small founder population colonizing both regions. However, in the within-haplogroup networks, the Faroe Islands are found within the larger set of potential source populations while Iceland is consistently found on isolated branches. Moreover, comparisons of within-haplogroup MDM histograms document a clear founder signal in the Faroes and Iceland, but the strength of this signal is haplogroup-dependent which may be indicative of more recent admixture or other demographic processes. The results of the current study and lack of conformity between Icelandic and Faroese haplotypes implies that the two populations were founded by different paternal gene pools and there is no detectable post-founder admixture between the two groups.

## 1 Introduction

> *There was a man named Grim Camban. He first settled the Færeys in the days of Harold Fairhair. For before the king’s overbearing many men fled in those days. Some settled in the Færeys and began to dwell there, and some sought to other waste lands. Aud the deeply wealthy fared to Iceland, and on her way thither she came to the Færeys, and there she gave Olof the daughter of Thorstan the Red in marriage: whence is come the greatest lineage of the Færey–folk, whom they call the Gate-beards, that dwell in Eastrey (Powell, 1896).*

The Faroe Islands consist of an archipelago of 18 small islands, located in the North Atlantic, between South Norway, Iceland and Scotland (Fig. 1). As a result of their demographic history and relative geographic isolation, the Faroe Islands, along with other North Atlantic Island populations, are genetically homogenous as compared to mainland populations (Wilson et al., 2001; Helgason et al., 2003; Goodacre et al., 2005). Historical and archaeological sources report that the Faroe Islands were settled around 800 CE by Vikings largely from western Norway (Arge, 1991; Arge et al., 2005). However, a growing body of evidence suggests that the islands were settled prior to this time, possibly by Celtic monks or other persons originating from the British Isles (Magnusson, 2003). Carbon dating of peat moss and barley grains support two pre-Viking settlement phases, around 300-500 CE and 500-700 CE (Church et al., 2013). More recently, Curtin et al. (2021) detected sheep-DNA in archaeological sediments from 500 CE, and based on modern whole-genome data, Gislason (2023) estimated that the initial founding of the Faroe Islands occurred between 50-300 CE, potentially two to three centuries earlier than previously believed based on archaeological findings alone.

**Figure 1:**
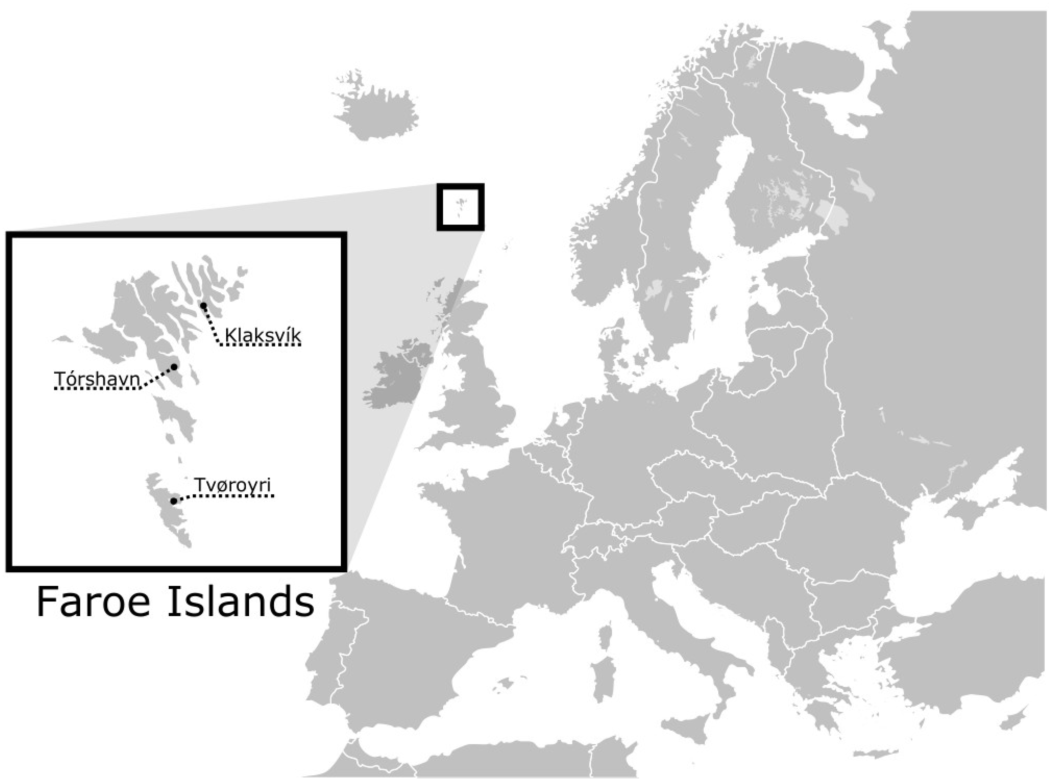
Geographic location of the Faroe Islands and the three cities where the samples were collected.

Genetic evidence suggests that, like other human populations on North Atlantic islands (including Iceland), the maternal and paternal founding populations of the Faroe islands are asymmetrical, with the vast majority of Faroese mitochondrial genomes being Celtic in origin, while most Y-chromosomal haplogroups are Scandinavian in origin (Jorgensen et al., 2004; Als et al., 2006). In addition, enslaved individuals from the British Isles of both sexes may have accompanied the early Faroes settlers and thralled Irish, Slavic, and Scandinavian individuals played an important role in early Viking society (Johnston, 1975; Jones, 1984). While it is unknown how large the founding population was, the demographic history of the Faroe Islands suggest that the islands were initially colonized by a small group of founders with a slow, but steady, increase in the population over time. During a 300 years period, from 1300 to 1600, the Faroese population was quite stable, around 3,000 to 4,000 people (Strøm, 2017). In 1600, it was estimated to be 3,200, but in 1650, it decreased to 2,515 individuals (Strøm, 2017). However, after 1700 there was a constant growth in the Faroese population; in 1700, around 4,000 people lived in the Faroes Islands, 5,255 in 1801, and 15,230 in 1901 (Strøm, 2017). In 2020, the population had reached 45,749 individuals. However, like other countries, the Faroe Islands have also been affected by different epidemics, which has caused the population to fall. During the years 1720 to 1865, the annual mortality rate (death per 1,000 inhabitants) fluctuated between seven and 42 (Strøm, 2017) (See Fig. S1 for more information).

To identify signatures of founder events in the Faroese population, we analyzed 12 individual Y-chromosomal microsatellite loci from 139 Faroese males (Table 1) and classified each haplotype into haplogroups using a Bayesian classification method (Athey, 2006). As Y-chromosomal haplogroups are defined by relatively slowly mutating single nucleotide polymorphisms (SNPs) and Y-chromosomal haplotypes are defined by relatively quickly mutating microsatellites, Short Tandem Repeat (STRs) (Zhivotovsky et al., 2004; Xue et al., 2009), we expect that haplotype diversity will recover faster than haplogroup diversity after a founding event. The fact that the vast majority of the Y chromosome is not subject to recombination indicates that hierarchical analysis of quickly mutating loci within groups delineated by slowly mutating loci may elucidate certain observations (de Knijff, 2000). Specifically, by definition, the occurrence of any SNP on the Y chromosome generates a novel haplogroup, the STR haplotype of which is singular, effectively resettling STR variation to zero. If the haplogroup is maintained in a population and increases in frequency, in subsequent generations haplotype variability will accumulate; in the absence of excessively high STR mutation rates, the original haplotype can be reliably inferred from modal values (de Knijff, 2000).

**Table 1:** Probability that a mutation will occur in any given generation (μ) for each locus typed in this study.

| Locus | $\mu$ | Reference |
| --- | --- | --- |
| DYS19 | 0.25% | NIST 2012 |
| DYS385a | 0.21% | NIST 2012 |
| DYS385b | 0.21% | NIST 2012 |
| DYS388 | 0.10% | Ravid-Amir & Rosset, 2010 |
| DYS389I | 0.24% | NIST 2012 |
| DYS389b | 0.35% | NIST 2012 |
| DYS390 | 0.25% | NIST 2012 |
| DYS391 | 0.28% | NIST 2012 |
| DYS392 | 0.07% | NIST 2012 |
| DYS393 | 0.08% | NIST 2012 |
| DYS426 | 0.07% | Ravid-Amir & Rosset, 2010 |
| DYS438 | 0.07% | NIST 2012 |

This study undertakes to further document the genetic impact of the colonization of the Faroe Islands. By using an extended number of STR loci for increased resolution, and by analyzing haplotype diversity in the context of haplogroups, a more precise, nuanced, and accurate understanding may emerge that clearly differentiates the evolutionary impact of the origin processes on the respective populations and permits exclusion of substantial post-founding paternal gene flow between Iceland and the Faroe Islands.

## 2 Materials and Methods

### Sample collection, DNA extraction, and sequencing

Buccal cells were collected using sterile swabs in 2004 and 2005 from 139 Faroes males living in three geographically disparate locations within the island group: (1) the city of Klaksvik (n=41) in the northern islands, (2) Tórshavn (n=48) in the central islands, and (3) Tvøroyi (n=50) in the south (Fig. 1). DNA was isolated from swabbed buccal cell samples using a standard phenol-chloroform extraction protocol.

Extracted DNA from each sample was next scored for 12 different Y-chromosomal STR markers (DYS19, DYS385a, DYS385b, DYS388, DYS389I, DYS389II, DYS390, DYS391, DYS392, DYS393, DYS426 and DYS438) in two multiplexes as described in Quintana-Murci et al. (2004) with primers and PCR conditions given in Redd et al. (2002). DYS389b was calculated by subtracting DYS389I from DYS389II (Kayser et al., 1997). Since DYS385 is characterized by a duplication, the shorter allele was assigned to DYS385a and the longer to DYS385b within each individual haplotype (Pacheco et al., 2005). To find potential parental sources to the Faroese population, buccal swabs from 47 Norwegian and 36 Swedish males were collected for the current study and were prepared in an identical manner as described above. Additionally, previously published haplotype data from Iceland (n=181) (Helgason et al., 2000), Denmark (n=185) (Hallenberg et al., 2005), and Ireland (n=148) (Ballard et al., 2006) were used as additional source populations (Table S1). Individuals within the putative parental data sets that had non-discrete or missing values were not included for the purposes of compatibility with the Faroese samples.

### Haplogroup inference

Haplogroups were inferred from haplotype data using an allele frequency-based Bayesian method (Athey, 2006), using nomenclature for haplogroups as defined by the Y-Chromosome Consortium (YCC) (Karafet et al., 2008). The online haplogroup prediction module (http://hprg.com/hapest5) allows the user to input specific marker values and returns a goodness of fit and probability score assigning an individual haplotype to one of ten common European haplogroups. In cases where individual haplotypes did not fall into expected Scandinavian or Celtic haplogroups (i.e., not R1a, R1b, or I1: n = 74 haplotypes), they were compared to the Y-Chromosome Haplotype Reference Database (YHRD) (https://yhrd.org) (Willuweit and Roewer, 2015).

Reliability of the Athey method of haplogroup inference was assessed by comparing Norwegian and Swedish samples that had been SNP-typed for I-P38 (I1*), I-P37b (I1b), I-P40 (I1a1), J-12f2a (J*), J-M172 (J2), N-tat (N3), R-M173 (R1*), R-P25 (R1b), R-SRY10831b (R1a) (The Y Chromosome Consortium, 2002). All Swedish samples were placed in the correct haplogroup. All Norwegian samples were also scored correctly save one SNP-tested R1a haplotype which scored a 62.3% probability for R1b and a 37.3% probability for R1a. Given its previous SNP testing, this sample was grouped with other R1a haplotypes for downstream analysis. Of all unique haplotypes included in this study (n=535) only 5% had a probability score of less than 90% (n=27). STR loci DYS388, DYS426, and DYS438 were only used for haplogroup inference, and are not included in subsequent analyses, as one or more were not reported in the complete Norwegian, Irish, Swedish, or Danish datasets.

### Haplotype network inference

Network analysis using high-resolution allele frequency data can parse distinct haplotypes within the same haplogroup (Zerjal et al., 2003; Moore et al., 2006). For each major haplogroup (R1a, R1b, I1), we generated haplotype networks using pairwise distances between all individuals (i.e., Hamming distance) generated with the R libraries adegenet (v2.1.10) (Jombart, 2008) and poppr (v2.9.5) (Kamvar et al., 2014). Correspondence analysis of all haplotypes across all haplogroups was performed with the R libraries adegenet (v2.1.10) (Jombart, 2008) and vegan (v2.6-4) (Oksanen et al., 2019). We calculated the degree of genetic differentiation among and between populations (Phi-statistics) within our haplotype networks using poppr::amova and the randtest function in the ade4 (v1.7-22) R library (Dray and Dufour, 2007).

### Histograms of within-haplogroup mutational distances

Assuming a stepwise mutation model (Kimura and Ohta, 1978), we estimated mutational distances from the modal haplotype (MDM) for each haplogroup. We define the MDM as the sum of the absolute number of repeat differences at each locus that an individual’s haplotype deviates from the population-specific modal haplotype which itself is determined by the modal repeat count at each locus. The MDM for each individual is therefore calculated as: 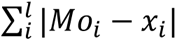

Where *Mo_i_* is the mode for any given locus within a haplogroup, *l* is the total number of loci, and *x* is the haplotype of an individual at locus *i*. This approach is an empirical extension of the method described by Rogers and Harpending (1992) who used mitochondrial data to calculate mismatch distributions.

Mismatch distributions of pairwise sequence differences have been shown to reproduce the structure of phylogenetic trees in simulated data (Slatkin and Hudson, 1991), and have been used to document expansions of human populations (Di Rienzo and Wilson, 1991). Specifically, the overall shape of the mismatch distribution and the position of the central tendency on the abscissa reflects time-since-expansion (Rogers and Harpending, 1992). This observation accords with the finding that in an exponentially growing population the expected phylogeny of haplotypes will be star-like and close to the coalescent (Slatkin and Hudson, 1991). We used the R moments library (v0.14.1) (Komsta and Novomestky, 2022) to quantify the degree of skewness (γ1) for each histogram wherein zero indicates complete normality, a positive value indicates a positive skew, and a negative value a negative skew. While empirical mismatch distributions do not mirror simulated populations at equilibrium, they reflect those of recently expanded non-equilibrium populations with reasonable accuracy (Rogers and Harpending, 1992). As our MDM histograms are generated using a modified pairwise comparison procedure, they should document similar distributional expectations with regard to time-since-founding and changes in demography. Given our interest in detecting a loss of diversity within haplogroups, comparing haplotypes to a locus-specific mode is a more appropriate methodology for our purposes.

We expect our MDM histogram analysis to represent the accumulation of within-population microsatellite diversity on a given haplogroup background since the time of the founding event. In the context of the expansion of an island population, predominant haplotypes in the modern population likely represent early founders. Consider a case where the small number of earliest founding males are paternally related; all will share identical (or near identical) Y-chromosomal haplotypes. During subsequent demographic expansion, the accumulation of mutations on founding lineages will be reflected in a progressive loss of zero mutation step instances in MDM histograms resulting in a gradual progression of the central tendency to the right.

All analysis scripts are available at https://github.com/aemann01/faroes_y. Scripts to calculate population specific modal haplotypes and generate MDM histograms are available as an interactive Shiny app at https://aemann01.shinyapps.io/mdmhistogram/.

### Diversity estimates

Finally, we calculated metrics of genotypic diversity, richness, and heterozygosity using the R poppr (v 2.9.5) (Kamvar et al., 2014) package. We reported four major diversity statistics across all haplogroups in all populations as well as haplogroup specific diversity metrics. Reported metrics include (1) MLG: the number of multilocus genotypes in each population, (2) eMLG: the expected number of MLG with a minimum of 10 samples after rarefaction, (3) lambda: a sample size corrected Simpson’s index (Simpson, 1949) calculated as: ((𝑁/(𝑁 − *1*)) ∗ 𝑙𝑎𝑚𝑏𝑑𝑎) where N equals the number of individuals in the population and lambda is the uncorrected Simpson’s index value, and (4) heterozygosity (Nei, 1978).

## 3 Results

### Faroese Y-chromosomal haplogroups

The most common haplogroups in all samples investigated here are haplogroups R1a, R1b, and I1 (Fig. 2a). Among the Faroese sample, these three haplogroups constitute 42%, 25%, and 21% of the dataset, respectively (Fig. 2a). Rare haplogroups were also found at low frequency. For example, haplogroup J1 constituted 4% of all Faroese samples, Q 3%, and haplogroups E1b1b, I2b1, I2b(xI2b1), I2a(xI2a1), L, and N combined make up approximately 6% of all Faroese samples (Table 2). Rare haplogroups found in source populations include G2a, J2b, J2a1b, and I2a1. Of the rare haplogroups found among the Faroese, six were found in one or more assumed source populations. The Faroese haplogroups J1 and L were not found in any source population. There is some inter-island variability between the Faroese sample populations, such that while haplogroup Q is shared among Tvorøyri, Klaksvík, and Tórshavn, haplogroups E1b1b, I2b, and I2a are found only on the northern island of Klaksvík, and haplogroups I2b1, L, and N are found only in the capital city of Tórshavn (Table 2). However, most of these haplogroups are represented by single individuals.

**Figure 2:**
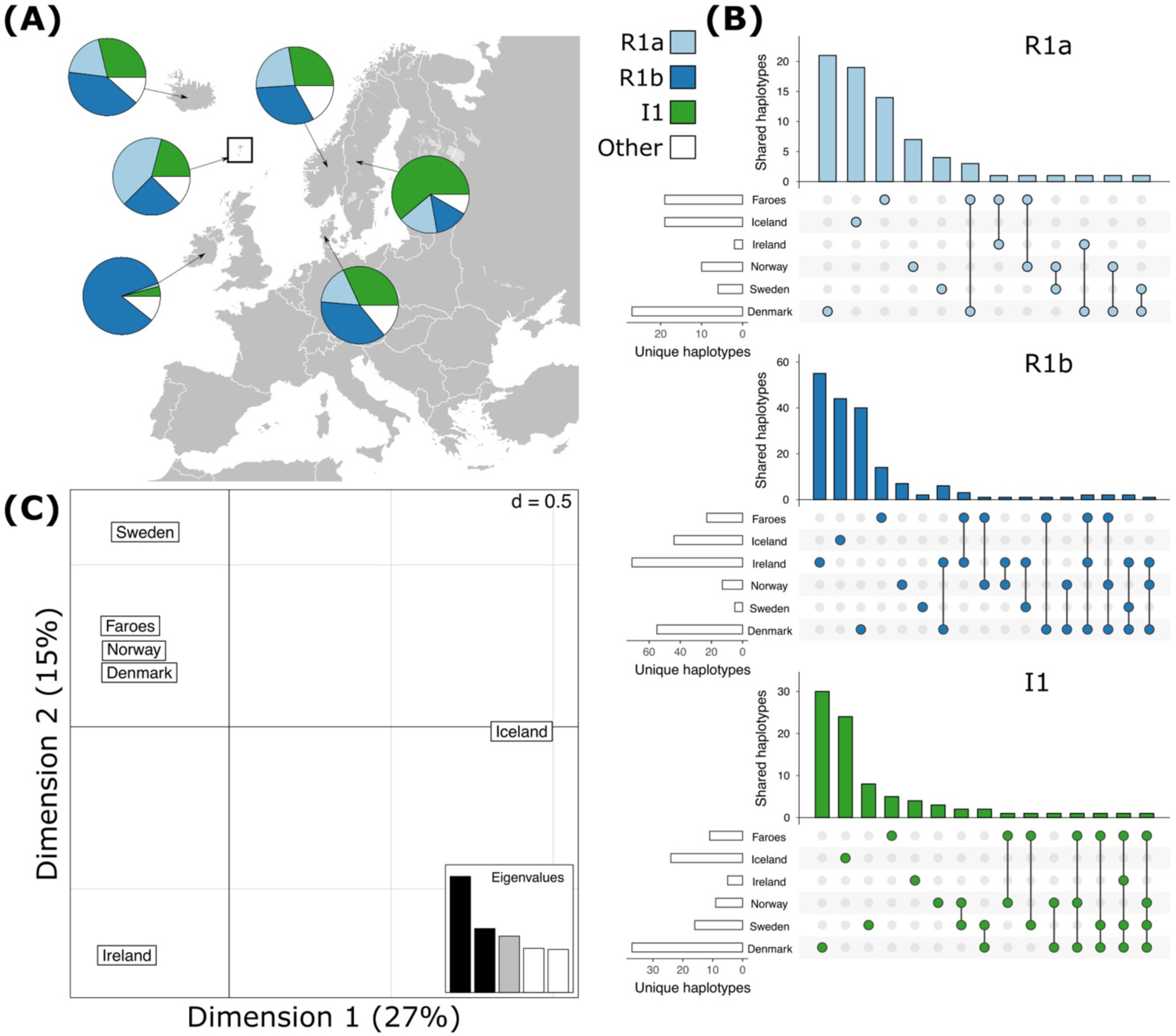
Haplogroup frequencies of the most common haplogroups, R1a, R1b, and I1 and shared haplotypes within haplogroups across all populations. (a) The most common haplogroups detected in each population. (b) Shared haplotypes (c) Correspondence analysis of all haplotypes across haplogroups for each population.

**Table 2:** Frequencies of different haplogroups in people living in Scandinavia and Ireland.

| Population | R1a | R1b | I1 | J1 | Q | E1b1b | I2b1 | I2b | I2a | L | N | Other * |
| --- | --- | --- | --- | --- | --- | --- | --- | --- | --- | --- | --- | --- |
| Faroese | 0.42 | 0.25 | 0.21 | 0.04 | 0.03 | 0.01 | 0.01 | 0.01 | 0.01 | 0.01 | 0.01 | 0.00 |
| Klaksvík | 0.37 | 0.27 | 0.15 | 0.10 | 0.02 | 0.05 | 0.00 | 0.02 | 0.02 | 0.00 | 0.00 | 0.00 |
| Tórshavn | 0.42 | 0.23 | 0.25 | 0.00 | 0.04 | 0.00 | 0.02 | 0.00 | 0.00 | 0.02 | 0.02 | 0.00 |
| Tvøroyri | 0.48 | 0.24 | 0.22 | 0.04 | 0.02 | 0.00 | 0.00 | 0.00 | 0.00 | 0.00 | 0.00 | 0.00 |
| Denmark | 0.17 | 0.37 | 0.32 | 0.00 | 0.02 | 0.02 | 0.05 | 0.01 | 0.01 | 0.00 | 0.01 | 0.03 |
| Iceland | 0.19 | 0.40 | 0.29 | 0.00 | 0.00 | 0.07 | 0.02 | 0.00 | 0.00 | 0.00 | 0.00 | 0.03 |
| Ireland | 0.01 | 0.84 | 0.04 | 0.00 | 0.00 | 0.01 | 0.05 | 0.00 | 0.03 | 0.00 | 0.00 | 0.02 |
| Norway | 0.23 | 0.32 | 0.28 | 0.00 | 0.02 | 0.00 | 0.06 | 0.00 | 0.00 | 0.00 | 0.06 | 0.02 |
| Sweden | 0.17 | 0.14 | 0.61 | 0.00 | 0.03 | 0.00 | 0.03 | 0.00 | 0.00 | 0.00 | 0.03 | 0.00 |

\*: Other haplogroups in **Denmark** include: G2a, J2a1b, J2b. **Iceland**: G2a, **Ireland**: G2a, J2b. **Norway**: J2a1h.

### Faroese Y-chromosomal haplotypes

Like haplogroups, haplotypes tend to be more variable among mainland Scandinavian samples than in the Faroese and Icelandic samples. Across all haplogroups, the lowest levels of genotypic richness (eMLG) and diversity (λ) are found among the Faroese (eMLG: 25.20, λ: 0.97), Icelandic (eMLG: 28.40, λ: 0.98), and Irish (eMLG: 29.70, λ: 0.98) populations but the levels of expected heterozygosity (Hexp) across all populations is fairly uniform with the lowest Hexp found among the Irish population (0.43) and highest among the Norwegian population (0.57) (Table 3). Percentages of unique haplotypes are highest among the Danish (88%), Norse (85%), and Swedish (83%) samples. The highest proportion of private haplotypes were in the Danish (59%), Icelandic (50%), and Irish (49%) samples. The Faroese samples are the least diverse at the haplotype level, such that 49% of the Faroese haplotypes are unique, while private haplotypes constitute 30% of the sample. Of the total unique haplotypes found in the Faroese sample, 31% (21 haplotypes) are shared with one or more source populations. The highest number of haplotypes assigned to R1a were shared among the Faroese and Danish populations followed by the Faroes and Ireland, and the Faroes and Norway (Fig. 2b). Among haplotypes assigned to R1b, the highest number were shared among the Faroes and Ireland, followed by the Faroes and Norway, and the Faroes and Denmark (Fig. 2b). Finally, among haplotypes assigned to haplogroup I1, the Faroes shared the most haplotypes with Norway, followed by Sweden (Fig. 2b). Considering all haplotypes independent of haplogroup identity, the Faroes cluster closely with Norway, Denmark, and to a lesser extent, Sweden using correspondence analysis (Fig. 2c).

**Table 3:** Total Estimates of Diversity. Estimates of diversity across all haplotypes independent of haplogroup assignment for each population. N indicates the number of individuals. MLG is the number of multilocus genotypes in the population. eMGL is the number of expected multilocus genotypes (i.e., genotypic richness) given the smallest sample size. 𝜆 is Simpson’s index (genotypic diversity) after controlling for sample size. Hexp is heterozygosity. k indicates the number of unique haplotypes found in a population. Percent unique is the proportion of haplotypes that are unique to the population (k/N). Percent private haplotypes is the proportion of haplotypes found solely in that population.

| Population | N | MLG | eMLG | $\lambda$ | Hexp | k | %unique | %private |
| --- | --- | --- | --- | --- | --- | --- | --- | --- |
| Faroese | 139 | 64 | 25.20 | 0.97 | 0.50 | 68 | 48.9 | 30.2 |
| Denmark | 185 | 144 | 33.70 | 1.00 | 0.55 | 163 | 88.1 | 58.9 |
| Ireland | 148 | 92 | 29.70 | 0.98 | 0.43 | 100 | 67.6 | 49.3 |
| Iceland | 181 | 98 | 28.40 | 0.98 | 0.51 | 120 | 66.3 | 49.7 |
| Norway | 47 | 40 | 31.60 | 0.99 | 0.57 | 40 | 85.1 | 48.7 |
| Sweden | 36 | 30 | 30.00 | 0.99 | 0.48 | 30 | 83.3 | 47.2 |
| Total | 736 | 468 | 34.20 | 1.00 | 0.62 | 521 |  |  |

#### Within–haplogroup haplotype diversity

Unlike estimates of expected heterozygosity that do not consider haplogroup membership, within-haplogroup haplotype diversities vary substantially depending on the haplogroup in question. For example, while the Faroese and Icelandic populations have the lowest within-haplogroup heterozygosity score (excluding populations with fewer than five representative individuals) for R1a (Faroes: 0.23, Iceland: 0.28) and I1 (Faroes: 0.17, Iceland: 0.20) haplogroups, within-haplogroup haplotype diversity metrics among R1b individuals are comparable across all populations (Table 4) with the lowest expected heterozygosity found among Norwegian haplotypes assigned to R1b (0.28) followed by the Faroese (0.32), Irish (0.34), Icelandic (0.35), Danish (0.36) and Swedish (0.40) haplotypes. Importantly, the most distinct and consistent differences between within-haplotype heterozygosity (Table 4) and total heterozygosity (Table 3) can be found among the Faroese and Icelandic populations. The low diversity observed among the Faroese and Icelandic populations is also reflected in the networks and histograms of mutational distances presented below.

**Table 4:** Within-Haplogroup Estimates of Diversity. Estimates of diversity across all haplotypes within one of the three most common haplogroups. N indicates the number of individuals. MLG *is the number of multilocus genotypes in the population. eMGL is the number of expected multilocus genotypes (i.e., genotypic richness) given the smallest sample size. 𝜆 is Simpson’s index (genotypic diversity) after controlling for sample size. Hexp is heterozygosity*.

| R1a Haplotype Diversity |  |  |  |  |  |
| --- | --- | --- | --- | --- | --- |
| Population | N | MLG | eMLG | $\lambda$ | Hexp |
| Faroese | 58 | 19 | 6.78 | 0.89 | 0.23 |
| Denmark | 31 | 27 | 9.54 | 0.99 | 0.37 |
| Ireland | 2 | 2 | 2.00 | 1.00 | 0.18* |
| Iceland | 35 | 19 | 8.03 | 0.95 | 0.28 |
| Norway | 11 | 10 | 9.18 | 0.98 | 0.40 |
| Sweden | 6 | 6 | 6.00 | 1.00 | 0.42 |
| Total | 143 | 83 | 9.18 | 0.98 | 0.40 |
| R1b Haplotype Diversity |  |  |  |  |  |
| Population | N | MLG | eMLG | $\lambda$ | Hexp |
| Faroese | 35 | 23 | 8.29 | 0.94 | 0.32 |
| Denmark | 69 | 69 | 9.62 | 0.99 | 0.36 |
| Ireland | 124 | 124 | 9.17 | 0.98 | 0.34 |
| Iceland | 73 | 73 | 8.95 | 0.97 | 0.35 |
| Norway | 15 | 15 | 9.14 | 0.98 | 0.28 |
| Sweden | 5 | 5 | 5.00 | 1.00 | 0.40 |
| Total | 321 | 321 | 9.76 | 0.99 | 0.45 |
| I1 Haplotype Diversity |  |  |  |  |  |
| Population | N | MLG | eMLG | $\lambda$ | Hexp |
| Faroese | 29 | 11 | 6.07 | 0.86 | 0.17 |
| Denmark | 59 | 37 | 8.98 | 0.97 | 0.31 |
| Ireland | 6 | 5 | 5 | 0.93 | 0.29 |
| Iceland | 52 | 24 | 6.48 | 0.83 | 0.20 |
| Norway | 13 | 9 | 7.38 | 0.92 | 0.25 |
| Sweden | 22 | 16 | 8.53 | 0.96 | 0.29 |
| Total | 181 | 102 | 9.21 | 0.98 | 0.40 |

### Identification of founder effect in haplotype networks

In the R1b haplogroup network (Fig. 3), several Faroese haplotypes stem from one larger common node, with other haplotypes dispersed across the network. The R1b network is dominated by the Irish sample with no population exclusively marking a cluster except for Iceland. Faroese haplotypes are somewhat centrally clustered in the network, but otherwise are highly dispersed throughout. Within the R1b haplogroup, Faroese haplotypes are shared with those found in Ireland, Norway, and Denmark (Fig. 2b) but never with Sweden or Iceland. Of the haplotypes from source populations, Danish haplotypes are widely distributed throughout the network – with the exception of the Icelandic branch. Danish haplotypes have the highest within population distance of all source populations with a mean of 4.0 (± 1.55). Mean genetic distance is also high among Swedish haplotypes (4.4, ± 1.65) but this is largely due to the small sample size of Swedish haplotypes assigned to this haplogroup (n = 5). For all other populations, the mean genetic distance is comparable: Faroes (3.54 ± 1.71), Ireland (3.76 ± 1.65), Iceland (3.79 ± 1.82), and Norway (3.04 ± 1.41). However, Faroese haplotype diversity within the R1b haplogroup (0.94, Table 3) is higher than that found in R1a and I1, implying that R1b haplotypes may have been more diverse in the original founding population. Alternatively, post-founding gene flow may have contributed to this higher degree of diversity (i.e., post-founding admixture with mainland Scandinavian populations). Among all major haplogroups, the degree of genetic differentiation between populations, while significant (p = 0.01) is lowest among R1b haplogroups (ΦPT = 0.29) and the majority of observed genetic variation is within populations (ΦPT = 0.71).

**Figure 3:**
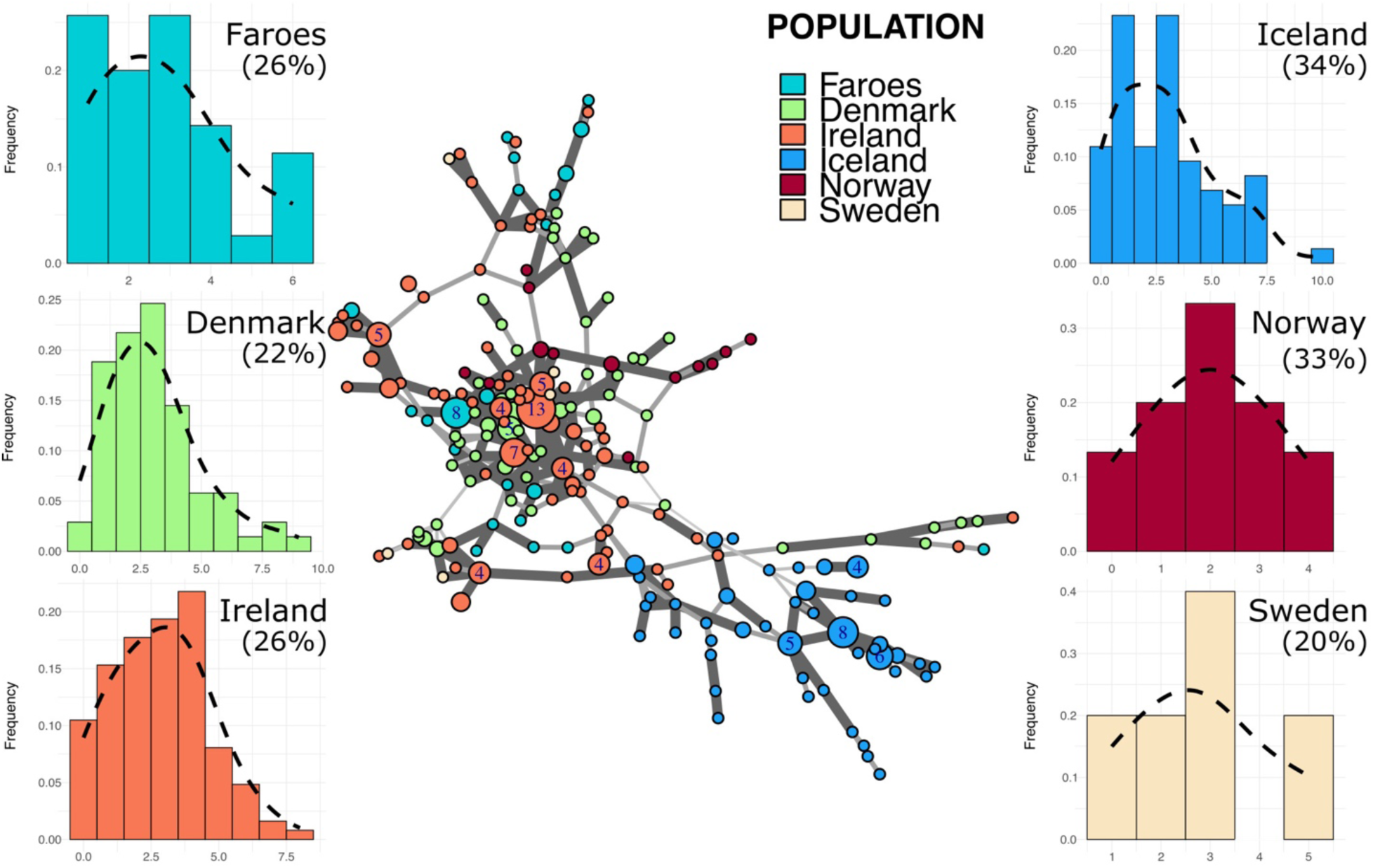
Minimum spanning network and mutational distance histograms of all R1b individuals. Minimum spanning network in the center of figure represents all haplotypes assigned to haplogroup R1b across all populations in this study. Color of node indicates source population while size of node represents the number of individuals assigned to that node. Larger nodes have numbers representing the exact number of individuals for that node. Thickness and opacity of lines connecting nodes correspond to the number of mutational changes (i.e., Hamming distance) between connected haplotype nodes where thicker, darker lines indicate fewer changes between nodes. Mutational distance histograms (MDM) for each source population illustrate the mutational distance for each haplotype from a population-specific modal haplotype. Percentages indicate the total proportion of haplotypes within each population that are “neighbor haplotypes”, i.e., haplotypes with only one repeat difference from the modal haplotype.

The R1a haplogroup network (Fig. 4) is substantially sparser as compared to the R1b haplogroup, primarily due to the lack of representation of our Irish population within this group (n = 2). In the R1a haplogroup network, the Faroe Islands are characterized by several large nodes that primarily cluster with Danish haplotypes with a smaller and more distinct node that clusters with Norwegian R1a haplotypes.

**Figure 4:**
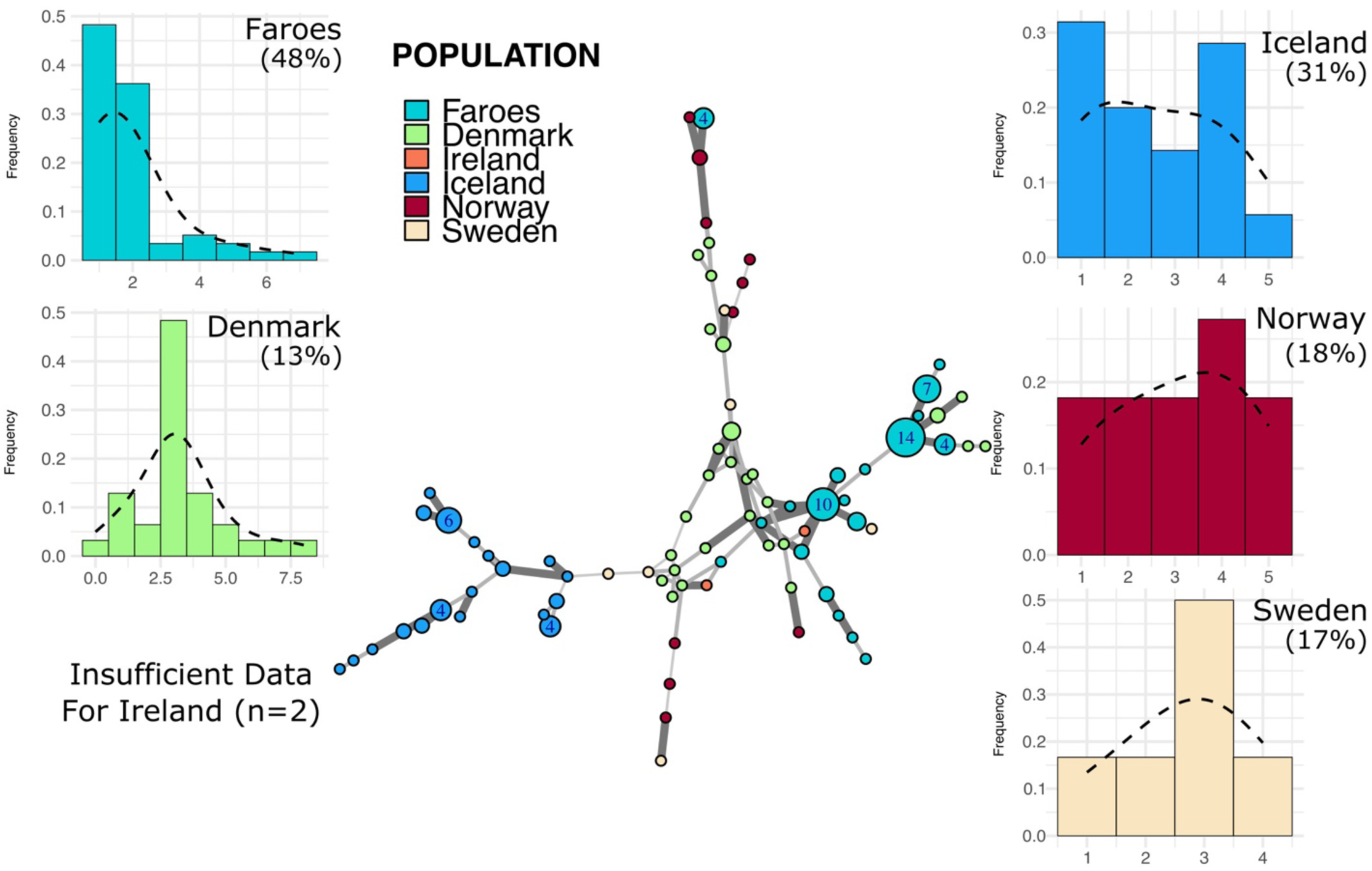
Minimum spanning network and mutational distance histograms of all R1a individuals. Minimum spanning network in the center of figure represents all haplotypes assigned to haplogroup R1a across all populations in this study. Color of node indicates source population while size of node represents the number of individuals assigned to that node. Larger nodes have numbers representing the exact number of individuals for that node. Thickness and opacity of lines connecting nodes correspond to the number of mutational changes (i.e., Hamming distance) between connected haplotype nodes where thicker, darker lines indicate fewer changes between nodes. Mutational distance histograms (MDM) for each source population illustrate the mutational distance for each haplotype from a population-specific modal haplotype. Percentages indicate the total proportion of haplotypes within each population that are “neighbor haplotypes”, i.e., haplotypes with only one repeat difference from the modal haplotype.

Like with haplogroup R1b, Faroese R1a haplotypes are shared with Denmark, Ireland, and Norway, but not with Sweden or Iceland. The mean genetic distance among all Faroese R1a haplotypes was substantially lower than any other source population (2.55 ± 1.62). Swedish R1a haplotypes had the highest mean genetic distance (4.6 ± 0.99) followed by Norway (4.35 ± 1.65), Denmark (4.04 ± 1.51) , and Iceland (3.08 ± 1.35). Too few individuals were assigned to this haplogroup in Ireland to perform mean distance analysis (n = 2). Faroese haplotype diversity within haplogroups is lowest in the R1a group (0.89, Table 3). Genetic differentiation among individuals is higher among R1a haplotypes (ΦPT = 0.34, p = 0.01) than R1b haplotypes but most of the genetic variation is similarly found within populations (ΦPT = 0.66).

Most Faroese haplotypes in the I1 network (Fig. 5) are closely clustered with two larger nodes and several smaller nodes that all fall central to other source populations with the exception, again, of Iceland. Faroese I1 haplotypes are shared with all source populations with the exception of Iceland (Fig. 2b). In addition to having somewhat lower haplotype diversity within the I1 haplogroup (0.86, Table 3), the Faroese I1 haplotype have the lowest mean genetic distance among all populations (1.83 ± 1.36) followed by Iceland (2.18 ± 1.58), Norway (2.79 ± 1.51), Ireland (3.13 ± 1.41), Sweden (3.23 ± 1.63), and Denmark (3.42 ± 1.7). We detected the highest degree of population differentiation among I1 haplotypes wherein 44% (ΦPT = 0.44, p = 0.01) of all genetic variation was found between populations and only 56% (ΦPT = 0.56) within populations.

**Figure 5:**
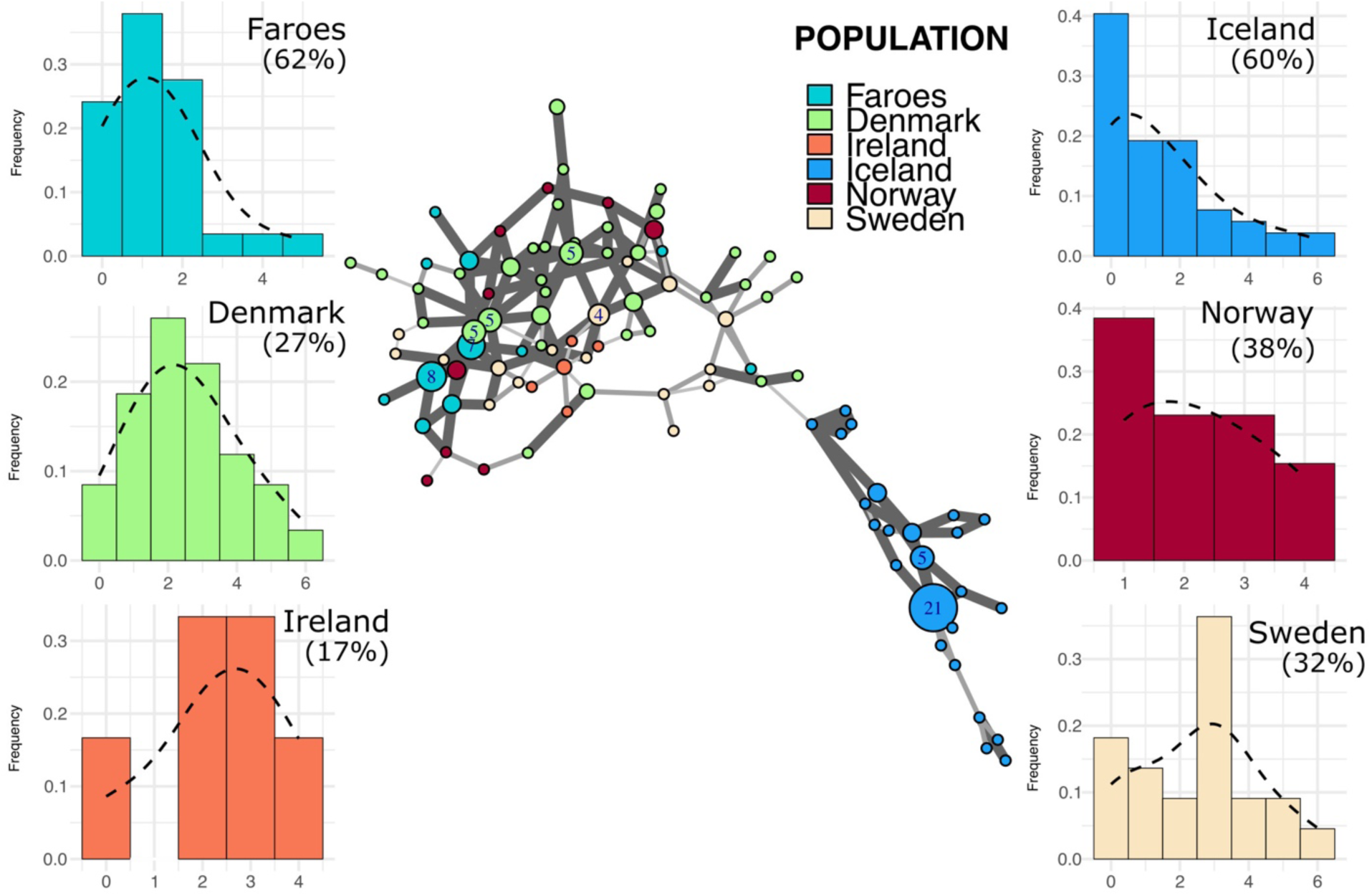
Minimum spanning network and mutational distance histograms of all I1 individuals. Minimum spanning network in the center of figure represents all haplotypes assigned to haplogroup I1 across all populations in this study. Color of node indicates source population while size of node represents the number of individuals assigned to that node. Larger nodes have numbers representing the exact number of individuals for that node. Thickness and opacity of lines connecting nodes correspond to the number of mutational changes (i.e., Hamming distance) between connected haplotype nodes where thicker, darker lines indicate fewer changes between nodes. Mutational distance histograms (MDM) for each source population illustrate the mutational distance for each haplotype from a population-specific modal haplotype. Percentages indicate the total proportion of haplotypes within each population that are “neighbor haplotypes”, i.e., haplotypes with only one repeat difference from the modal haplotype.

To summarize, the distribution of Faroese haplotypes across minimum spanning networks within haplogroups do not match the stereotypical patterns for recently founded populations; specifically, the presence of clusters of singleton haplotypes surrounding large central nodes in star-like formations.

However, we do find differences in population differentiation across major haplogroups with different degrees of observed genetic variation within and between groups, supporting the idea that within haplogroup diversity estimates may have finer resolution for documenting population events such as founder effect.

### Identification of founder effect in the distribution of mutational distances

Histograms representing stepwise mutational distances from within-haplogroup modal haplotypes for each population are found in Figs. 3-5. We hypothesize that different distribution patterns around modal haplotypes indicate various demographic processes involving population size, age, and the relative success of individual lineages. In the case of recent founder effect, we expect that the majority of mutational distances from the modal haplotype to be a single mutational step away or fewer (i.e., the modal haplotype) which we have termed “neighbor haplotypes”. In addition to neighbor haplotypes, we calculated the degree of skewness (γ1) for each histogram wherein a value of zero indicates complete normality, a positive value indicates a positive skew, and a negative value a negative skew. The results of this analysis reveals two basic patterns among all populations, mainly smooth distributions with a single mode and multi-modal distributions.

Putative source populations (i.e., Denmark, Ireland, Norway, and Sweden), generally have MDM distributions characterized by a single mode, but the mode has shifted further away from zero with a lower overall proportion of neighbor haplotypes, indicating the kind of accumulated variability expected in older populations. In contrast, the Faroese and Icelandic sample tends to have modes nearer to zero with a higher proportion of neighbor haplotypes and are skewed to the right. This pattern, however, is haplogroup specific which may reflect differences in the diversity of the original founding group, admixture post founding, or other aspects of population structure. For example, the distribution of haplotypes from the modal haplotype within haplogroup I1 is highly skewed towards the right among the Faroese (γ1 = 1.19) and Icelandic (γ1 = 1.13) populations with 60% or more haplotypes representing neighbors to the mode (Fig. 5). Similarly, Faroese haplotypes are strongly skewed to the right within the R1a haplogroup (γ1 = 2.02) with 48% of all haplotypes representing neighboring haplotypes.

Interestingly, while a relatively high proportion of Icelandic haplotypes within the R1a haplogroup are neighbors to the mode (31%), the distribution of Icelandic R1a haplotypes is multimodal (γ1 = 0.17) which reflects underlying population structure also seen in our minimum spanning networks. This underlying population structure is mostly clearly seen among MDM histograms representing R1b haplotypes where there is a clear multimodal distribution that is skewed towards the right among both the Faroese (γ1 = 0.63) and Icelandic (γ1 = 0.78) haplotypes but a smooth accumulation of mutational distance from the mode among Danish (γ1 = 1), Irish (γ1 = 0.28), Norwegian (γ1 = 0), and Swedish populations (γ1 = 0.37). To test whether these multimodal distributions found among the Faroes and Icelandic R1b and R1a haplotypes are the result of true within-haplogroup diversity or haplogroup classification error, we performed an additional “strict” haplogroup MDM histogram analysis where to be included a haplotype must have had a probability score above 90% and a fitness score of > 50 using the Athey haplogroup prediction program. Notably, however, the proportion of neighbor haplotypes among all populations are similar despite a clear shift from the modal haplotype among our North Atlantic Island populations. Interestingly, while the multimodal distribution previously observed among the Faroese and Icelandic R1a haplotypes diminished significantly using these strict parameters, the multimodal distribution found among the Faroese R1b haplogroup remained relatively unchanged (Fig. S2).

Despite variation among different haplogroups, these results follow the expectation that source populations have broader and smoother distributions, notwithstanding sample size and the effect of latent population structure. In contrast, the Faroese sample has modes nearer zero and are skewed to the right.

## 4 Discussion

The Vikings played a preeminent role in the peopling of the North Atlantic, and one might expect populations that were founded by the Vikings to be genetically similar and homogeneous. Previous investigation of Y–chromosomal haplotype and haplogroup frequencies in the Faroe Islands found a high degree of similarity between the Faroe Islands, Norway, Sweden, and Iceland, indicating a Scandinavian ancestry among the male settlers of the Faroe Islands (Jorgensen et al., 2004). Faroese mitochondrial lineages, however, indicates an excess of British Isles ancestry among the female settlers of the Faroe Islands (Als et al., 2006). In both studies the Faroese population was found to be highly homogeneous, reflecting an expected small founder population size and strong genetic drift. In the case of the Y– chromosome, there were relatively few haplotypes per sample, relatively low heterozygosity, and a correspondingly low effective population size (Jorgensen et al., 2004). In the current study we argue that the application of a moderately extended haplotype in conjunction with consideration of estimates of diversity in the context of network analysis can more clearly elucidate founder effect in the Faroese population. Specifically, we argue that the opportunity to detect loss of diversity associated with founder effect will be affected by the level of phylogenetic resolution considered. The significance of hierarchical analysis for this study is that estimates of haplotype diversity may be inflated by underlying haplogroup diversity and the degree to which those haplogroups have diverged in conjunction with their position on the Y–chromosomal gene tree. Thus, an improved assessment of hierarchical levels of diversity enhances our understanding of the genetic history of the Faroe Islands and the consequences of the founder event.

### Signals of founder effect

#### Private and unique haplotypes

The most constrained genetic diversity is consistently found in the Faroese samples. The sample from Faroe Islands has the lowest percentages of unique haplotypes (i.e., the total number of unique haplotypes within a population) as well as private haplotypes (i.e., haplotypes found in that population while completely absent from all other populations). Percentages of unique haplotypes within Scandinavian populations—Norway, Sweden, and Denmark—are considerably higher than those found in the North Atlantic Island populations (i.e., Faroe Islands and Iceland). Unique haplotypes are frequently found as a result of demographic expansion (Slatkin and Hudson, 1991). The low percentage of unique haplotypes in the Faroe Islands may thus reflect its demographic history.

Percentages of private haplotypes within the Scandinavian and Icelandic populations are also considerably higher than the Faroe Islands (Table 3); there is not such a straightforward explanation for this result. Low proportion of private haplotypes in contemporary the Faroese population is indicative of comparatively recent small–scale founder events with little evidence for a re–accumulation of haplotype diversity. Genetic drift in isolated populations with a small founding number decreases heterozygosity within the population and increases the differentiation between populations (Chiaroni et al., 2009).

The frequency of private haplotypes found in a given population can be influenced by the degree of sampling saturation, divergence from other populations, and genetic drift (Helgason et al., 2000). The presence of private haplotypes is also a function of genetic resolution, such that the more microsatellites scored, the more likely a given haplotype will be private within a population. Additionally, differences of private haplotypes may also be partly the consequence of sampling strategy. Population structure can be found in large, geographically dispersed source populations, and is best captured with broad geographic sampling (cf. Dupuy et al. (2006) for the example of Norway). In the present study, the broad geographic sampling scheme for Sweden may explain minimal sharing of haplotypes (6%), while Jorgensen et al. (2004) reported 34% of Faroese Y–chromosomes were assigned to Sweden.

#### Parsing haplotype diversity

Measurements of diversity will be affected by whether haplotype variability is considered without respect to haplogroup. Sampling without regard to the distribution of haplotypes on a haplogroup evolutionary network is analogous to sampling a large population without regard to population structure. The significance of this is that haplotype diversity estimates will be inflated depending on the degree of haplogroup diversity found in the population: Disregarding haplogroup identity will obfuscate a signal of founder effect. Each new haplogroup represents a different evolutionary history, and it is inappropriate to look at microsatellite diversity by combining data from across many different haplogroup lineages. In the present study, haplotype heterozygosity estimates from island populations are comparable to source populations when haplotypes are considered without respect to haplogroup, yet within–haplogroup heterozygosity estimates are substantially lower (Table 3).

Consider the fact that within–haplogroup haplotype diversity is low in the Faroese sample with the lowest haplotype variability found in the most frequent haplogroup, R1a (h=0.89); whereas haplotype diversity calculated without regard to haplogroup in the Faroe Islands is substantially higher (Total ĥ=0.97), thereby masking a signal of founder effect. This line of reasoning likely extends to other summary statistics.

### Signals of founder effect on networks

#### Faroese haplotypes are distributed on haplogroup networks

It is a general expectation that star–like clusters of singleton haplotypes crowded around large primary nodes in haplotype networks are concordant with founder events and subsequent expansion. This has been demonstrated in Y– chromosomal studies of Ashkenazi Jews (Nebel et al., 2005), the population living on the Indonesian island of Nias (van Oven et al., 2011), and in the Irish (Moore et al., 2006). In addition to some primary nodes, several Faroese haplotypes are found widely dispersed on the networks. The position and distribution of Faroese haplotypes across networks depends on certain variables: the effective population size of the initial founders in conjunction with the impact of genetic drift since the founding event; the relative contribution of more or less divergent haplotypes to the founding population; and the degree of reproductive isolation since the time of founding. If the original effective size was small, we would expect to see a few larger nodes with closely related haplotypes clustered nearby. If paternal contributions are geographically and phylogenetically diverse, Faroese haplotypes may show up scattered across each network. In the present study, we find generally that singleton Faroese haplotypes are widely distributed across each haplogroup network, the majority of which are found either in large clusters of primary haplotype nodes and shared with one or more source populations, or close one– and two–step neighbor haplotype nodes. The distributions of Faroese haplotypes across the haplogroup networks demonstrate that the Faroese colonizers originated from multiple source populations, while classic founder effect is demonstrated by low within–haplogroup diversities.

#### Icelandic haplotypes are isolated on networks

Although within–haplogroup haplotype diversity estimates in samples from the Faroe Islands and Iceland are comparatively lower than source populations, the degree of Icelandic haplotype isolation in each network is distinctive. For each of the haplogroup networks, most Icelandic haplotypes are tightly clustered to the exclusion of haplotypes from other populations; these isolated clusters of haplotypes are not shared with any source populations or the Faroe Islands. Both the Faroese and Icelandic founders are thought to have been mostly Vikings who originated primarily from Norway, Sweden, or Denmark, and the distribution of Faroese haplotypes on the networks demonstrates this relationship. That the isolated clusters of Icelandic haplotypes are not shared may be explained by incomplete sampling of source populations, such that the region(s) where Icelandic founding haplotypes originated was not sampled. A larger, more diverse set of source population samples with greater regional differentiation may clarify the respective origins of the Faroese and Icelandic founder groups. It is also plausible that the Icelandic founder population originated from long–established Viking colonies in the British Isles, while the Faroese are the descendants of a more direct Scandinavian settlement process. Regarding the initial Icelandic settlers, such a scenario was proposed by Helgason et al. (2000). Thus, successive and serial bottlenecking of genetic variability of the Icelanders coming by way of Ireland likely explains both their position and stark clustering in the haplogroup networks.

### Mutation distance from the modal haplotype histograms

Faroese and Icelandic R1a and I1 haplogroup MDM histograms are typical of our expectations for a signal of founder effect, that is, highly leptokurtic distributions which are skewed to the right. As the modal haplotype is constructed by calculating the mode for each locus and combining them, it is expected that, in a population recently founded, most individuals within the haplogroup will share the modal haplotype. These MDM histograms reveal a recent, small–scale founder event in that most haplotypes are typically less than two mutational steps away from the calculated mode. The Faroese pattern is distinct from that of the source populations where the central tendency of a given source population is often at two or more mutational steps from the calculated mode. The Icelandic distribution for R1a is slightly shifted to the right with a mode of two; this is best understood in light of the isolated collection of Icelandic haplotypes, which are moderately divergent from themselves.

These results contrast somewhat with those of haplogroup R1b. Bearing in mind that R1b is an older haplogroup than both R1a and I1 (Hallast et al., 2015), greater haplotype diversity is expected. In the case of the Faroe Islands, the most notable feature is a strong multimodal distribution with a relatively high number of haplotypes that are six mutational steps from the calculated mode. Similarly, Iceland is multimodal with a relatively high proportion of haplotypes that are highly divergent from the mode, a pattern similar to that seen in the Danish and Irish R1b haplotypes. It is possible that this pattern is the result of convergence during founding of two distinct branches contributing divergent haplotypes contributing to the original R1b founders of Iceland (cf. Helgason et al., 2000). Alternatively, these patterns may indicate more recent admixture from Denmark. The Faroe Islands are today a self-governing part of the Danish Realm. For centuries, however, there has been a close relationship between Faroe Islands and Denmark, mainly political and administrative, but also on family and personal levels. It is likely that this shared political and demographic history led to gene flow between the two countries.

### Variability between regions of the Faroe Islands

Three geographic locales were sampled in the Faroe Islands: the northern islands (Klaksvík), central islands (Tórshavn), and southern islands (Tvøroyri). Due to the geographic isolation of the Faroe Islands and high mobility of individuals within the island group, it is expected that collectively the locales will be genetically homogeneous. Although likely insignificant from the standpoint of population structure, historical contingencies and modern transport with cars and ferries has reduced the human isolation between the islands which has impacted the genetic variation within the Faroese as a whole.

Each region is characterized by a high frequency of R1a, R1b and I1 and the inclusion of rarer groups (Table 2, Fig. S3). Similar to previous genetic studies that examined Y–chromosomal haplotypes (Jorgensen et al., 2004), R1a is found to have the lowest amount of inner diversity in this study despite its high frequency among the Faroese peoples (42%): This high frequency and low diversity documents the effect of genetic drift. The homogeneity of this group is exemplified by the fact that most individuals (72%) are either part of or neighbor haplotypes to the modal node (Fig. 4). The higher frequencies of R1a among Western and Central modern Norwegian populations (32%) (Dupuy et al., 2006) are consistent with the premise that the original founder population of the Faroe Islands came from western Norway.

Although most of the Faroese population belongs to one of three highly frequent haplogroups, there are some rare haplogroups present in the population that are infrequent even among Europeans as a whole, and local frequencies of rare haplogroups vary between Faroese locales (Table 2). Minor structure at the haplogroup level may support an inference that while mobility between islands in the Faroe Islands is high, the northern islands are more genetically diverse than the central or southern islands, even though the geographic air distance between Klaksvík in the north and Tórshavn in the center is only around 25 km, and around 50 km between Tórshavn and Tvøroyri in the south. Future investigations using a more intensive sampling scheme may resolve the issue of island heterogeneity further. Regardless, this higher haplogroup diversity in the northern locale has implications for interpreting haplotype diversity between locales. The higher proportion of rare haplogroups found in the sample from Klaksvík (21%) is driving its relatively higher proportion of unique and private haplotypes, and indeed impacts the diversity estimates for the Faroe Islands as a whole.

Specific haplogroups found in this sample are indicative of migration after the initial founding of the Faroe Islands. This assertion is based on the expectation that haplogroups present in the initial founders should be at higher frequency. The only rare haplogroup of note is J1, which constitutes 10% of the Klaksvík Y chromosomes. Haplogroup J is thought to have originated in the Middle East and is associated with the origins and spread of early farming groups. Unlike the European J–M172 sub– haplogroup, however, J1 is most closely associated with Middle Eastern, northern African, and Ethiopian populations (Karafet et al., 2008). Haplogroup J1 is not found in any source population investigated in this study and is therefore highly unlikely to have originated in the founding group. Perhaps most intriguing is historical evidence that may account for this haplogroup’s presence in the Faroe Islands, specifically an attack by the Barbary corsairs in the 1600s (West and Jákupsson, 1980). Stories about Turkish pirates who came to the southernmost islands, Suðuroy, are well known among Faroese peoples. Modern populations living in regions where Barbary corsair ports once existed still possess high frequencies of the J1 haplogroup (Arredi et al., 2004). The J haplogroup is also found at higher than expected frequencies in other Atlantic island populations with complex colonization histories; an investigation into Y–chromosomal diversity in the Azores revealed that J haplogroup frequencies were twice those in mainland Portugal (Pacheco et al., 2005). Regardless of the source, haplogroups that are rare within the overall Faroe Islands sample inflate haplotype diversity when estimates are calculated without regard to haplogroup identity. This follows from the observation that haplotypes of regionally uncommon haplogroups will be more divergent from more common haplogroups and thus have a higher probability of being both private and unique.

### Faroese-Icelandic comparison

Although little is actually known about the settlement process of the Faroe Islands, it has largely been assumed that the settlement of the Faroe Islands was similar to that of Iceland (Arge et al., 2005), and historical data document the possibility that the Faroese and Icelanders share common founders. Previous genetic studies using lower resolution Y–chromosomal data have found close connections between the Faroese and Icelanders (Jorgensen et al., 2004). It has been estimated that Y–chromosomal lines of descent can be reliably discerned over 1950 generations (or 49,000 years) (de Knijff, 2000). Therefore, if Iceland and the Faroe Islands do share a common paternal source—or if there has been post–founder contact between the two populations—the data should reflect this at the haplotype level within major haplogroups.

Primary clusters of Icelandic haplotypes are always separate from the Faroese haplotypes within each major haplogroup. The minimum spanning networks of all populations present a conspicuous pattern of Icelandic haplotypes: most Icelandic haplotypes in all three major haplogroup networks consistently cluster on cohesive separate branches. This distinctive pattern of clusters is unique to that population. The overall Icelandic pattern stands in contrast to that of the Faroese. Faroese samples are fairly dispersed in all three networks. Large Faroese nodes tend to be located proximally within the network while private haplotypes are located more distally throughout, and none of the nodes are distinctly derived. These results stand in contrast to the distributions of the haplotypes from the source populations, which are much more dispersed. Taken together, these results provide little evidence to support the hypothesis that the Faroese and Icelanders originated from the same proximate source populations or have undergone subsequent episodes of admixture.

### Distinctive Icelandic founding fathers

The lack of obvious concordance of founders between the Faroe Islands and Iceland in the current study is unexpected, as a previous Y chromosome study found a high degree of similarity between the two populations (Jorgensen et al., 2004). This discrepancy is most likely due to the number of microsatellite loci used in the original study (five) and that of the current study (nine of the 12 loci presented here).

Divergence can be more easily detected if more Y microsatellite loci are available for comparison, and these loci are relatively more mutable. Four relatively mutable loci were used here that were not scored in the previous study (Table 1). The high level of clustering and isolation on the networks of the Icelandic samples is evocative of a latent SNP present in the sample. Because this study is confined to a particular level of haplogroup resolution, the representation of latent SNPs in the data can only be inferred and not implied directly. If the haplogroup defined by the latent SNP is sufficiently divergent from other haplogroup members and there are enough representative individuals within the data, clustering of a subgroup may be apparent within the larger haplogroup itself. This method has been used to distinguish sub–haplogroups in Northern Ireland (Moore et al., 2006) and particularly successful male lineages across Asia (Zerjal et al., 2003).

If Iceland and the Faroe Islands are indeed genetically divergent, as found in the present study, two scenarios may explain their differentiation and Iceland’s relative isolation on the networks: (1) the populations were settled by different regional groups; or, (2) another historically contingent process of evolution between the two groups has resulted in the divergence. In the case of the former, remote, geographically divergent Norwegian sources may have generated the multimodal distribution evident in R1b (Dupuy et al., 2006). In the case of the latter, as mentioned previously, and proposed by Helgason et al. (2000), multiple bottlenecks of genetic variation of Vikings settling in Ireland and those same Vikings colonized Iceland may also lead to isolation on networks. These scenarios are not mutually exclusive, and our data lack sufficient resolution to distinguish between these scenarios. Both scenarios can act to generate isolation on the network and the higher proportion of private haplotypes compared to the Faroe Islands sample. Both scenarios are consistent with reports of paucity of Icelandic genetic variation (as a function of genetic drift) (Helgason et al., 2003) and evidence for admixture incorporating divergent lineages from Scandinavia and Celtic populations giving rise to deeper heterogeneity even in the context of founder effect (Arnason, 2003; Helgason et al., 2000).

Regardless, our results suggest that the Icelandic and Faroese populations had distinguishably different founding fathers. Minimum spanning networks consistently place the Icelandic group apart from all other populations while the Faroese tend to be well dispersed. Data presented here also suggest that the Faroese male population was founded by a more diverse group from divergent Scandinavian populations than their Icelandic neighbors. Furthermore, we conclusively demonstrate that there is no evidence for post– founder admixture between the Faroese and Icelandic Y–chromosomal gene pools.

## 6 Author contributions

AEM: Investigation, Methodology, Visualization, Writing – original draft, Writing – review & editing; EM: Collection of samples, interpretation of data and writing and revision of the manuscript. CRT: Project administration, Methodology, Supervision, Conceptualization, Writing – original draft, Writing – review & editing

## 8 Acknowledgements

Many thanks to Tasha Altheide, Matt Kaplan, Arthur DeFruscio, and Neha Angal for fruitful discussions and suggestions for revision.

## 9 Data availability statement

The data analyzed for this study can be found in Supplementary Table 1. All analysis scripts can be found at https://github.com/aemann01/faroes_y and are archived at Zenodo under DOI: 10.5281/zenodo.12609965.

---

**Supplementary Figure 1:**
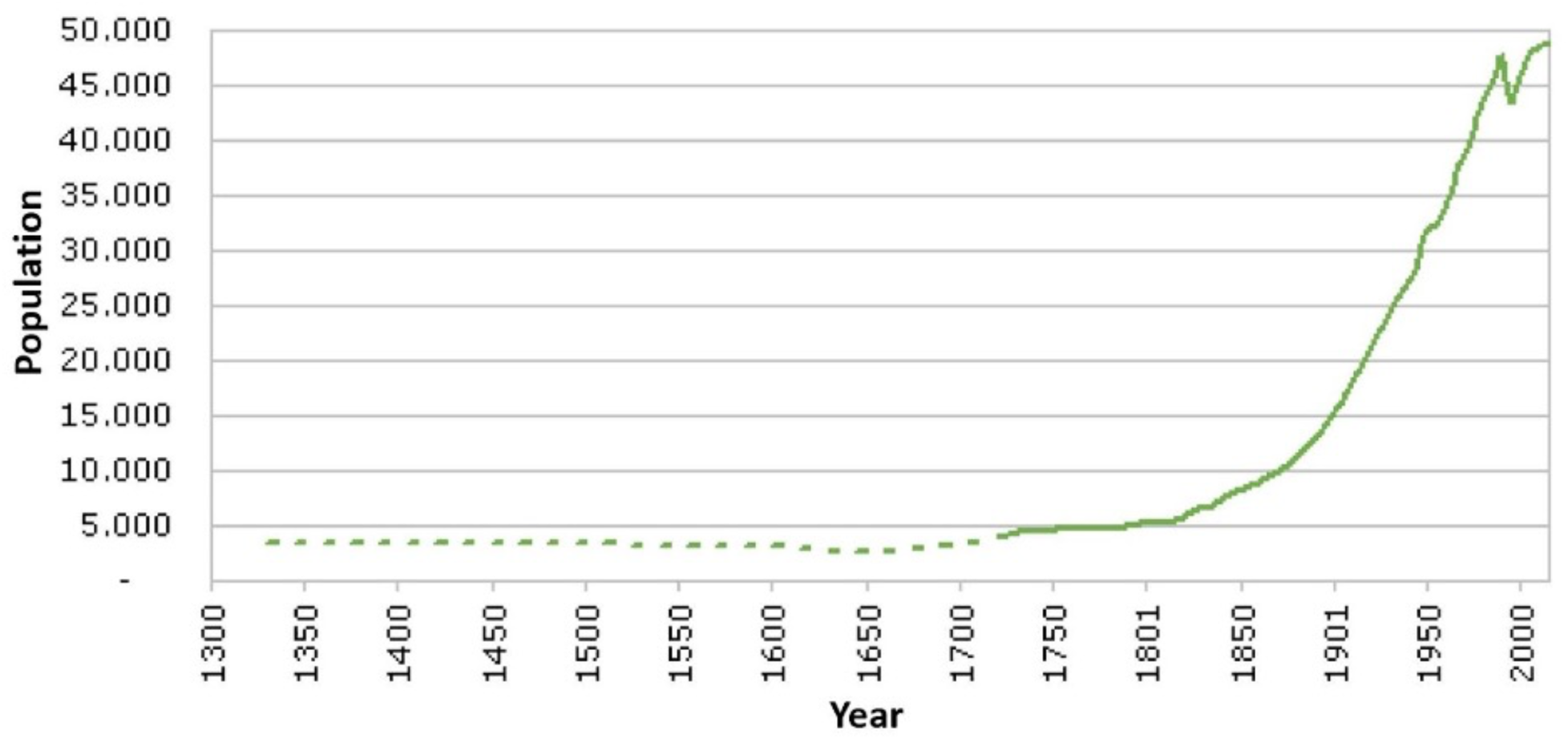
Population growth of the Faroese human population during 700 years, from 1330 to 2015. Data from Strøm 2017.

**Supplementary Figure 2:**
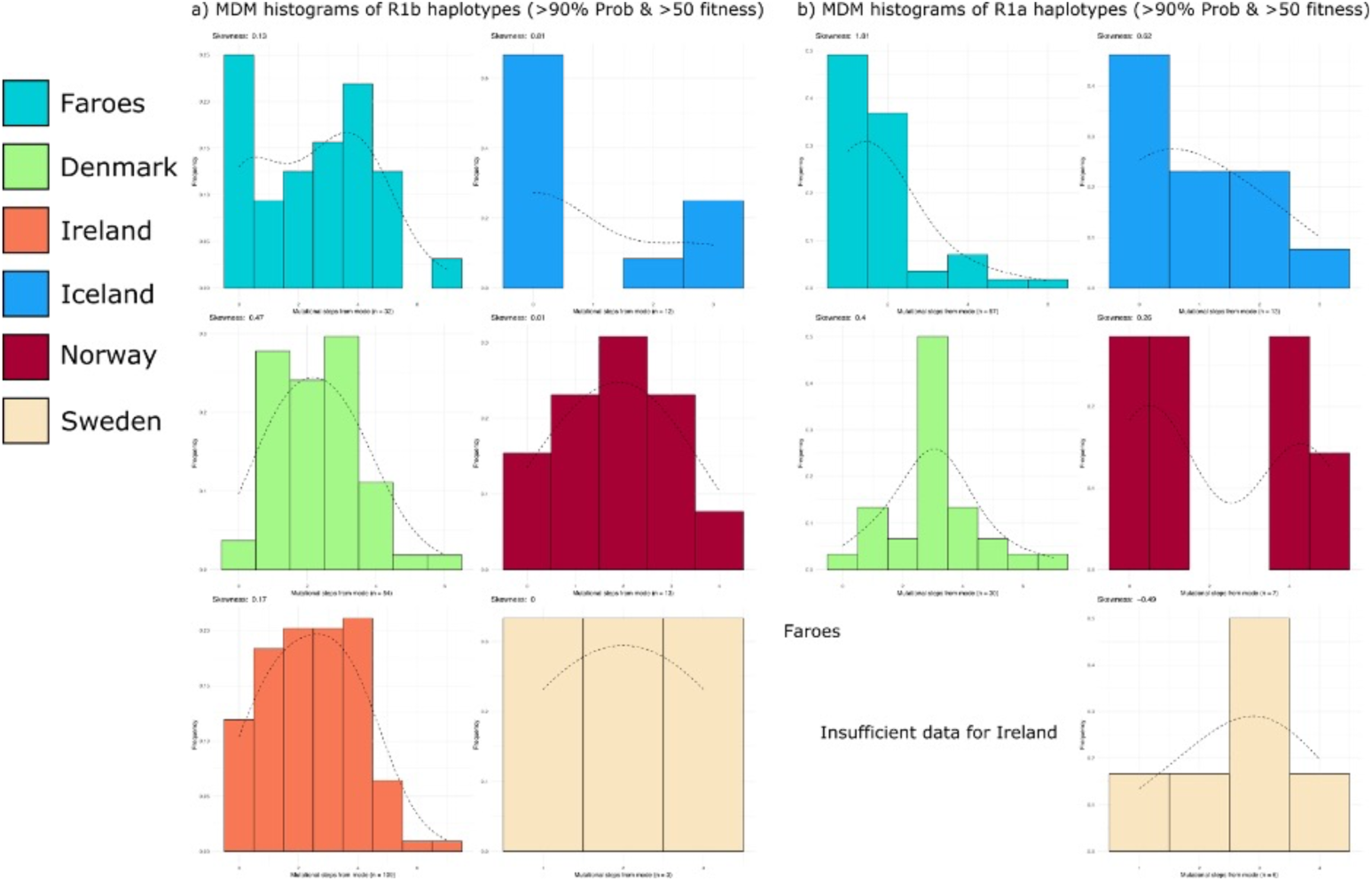
MDM histograms using strict haplogroup prediction. a) Population specific MDM histograms of haplotypes assigned to haplogroup R1b that passed strict haplogroup assignment (Probability > 90%, Fitness score > 50). b) Population specific MDM histograms of haplotypes assigned to haplogroup R1a that passed strict haplogroup assignment (Probability > 90%, Fitness score > 50).

**Supplementary Figure 3:**
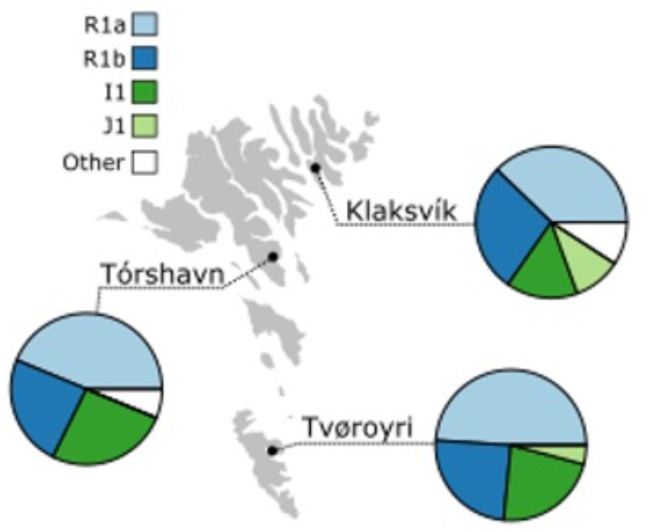
Haplogroup frequency by location in the Faroes. Haplogroups with less than 10% total frequency at any location collapsed into “other”.

